# Different strategies to manage *Varroa jacobsoni* and *Tropilaelaps mercedesae* in Papua New Guinea

**DOI:** 10.1101/2019.12.17.880393

**Authors:** John MK Roberts, Cooper N Schouten, Reuben W Sengere, John Jave, David Lloyd

## Abstract

Apiculture in the Pacific nation of Papua New Guinea (PNG) is under significant pressure from emerging parasitic mites, *Varroa jacobsoni* and *Tropilaelaps mercedesae*. While numerous mite control products exist, beekeepers in Papua New Guinea have limited access and resources for these products and their effectiveness under local conditions is untested. Here we determined the effectiveness of two brood manipulation strategies – queen caging and queen removal – for managing *V. jacobsoni* and *T. mercedesae* in comparison to the chemical miticide Bayvarol^®^. Our results found Bayvarol^®^ was the most effective control strategy for *V. jacobsoni*, maintaining high efficacy (>90%) over four months with significantly reduced mean numbers of *V. jacobsoni* compared to untreated control hives. In contrast, mean numbers of *T. mercedesae* were significantly reduced by the brood manipulation strategies over two months, but not by Bayvarol^®^ compared to the controls. These results highlight that a combination of different strategies is likely needed to effectively manage both mite pests in PNG. We discuss how these strategies could be optimised and adopted to achieve better mite management for PNG beekeepers.

## Introduction

The ectoparasitic mite, *Varroa destructor*, has become a major global pest of honey bees (*Apis mellifera*) after shifting hosts from the Asian honey bee *A. cerana* early last century (Anderson and Trueman, 2000). However, other serious mite threats are emerging in the Asia-Pacific region that have similarly shifted from their Asian honey bee hosts. *Tropilaelaps mercedesae* is an endemic parasite of the giant Asian honey bee (*A. dorsata*) that has since spread onto *A. mellifera* and become widespread throughout Asia where it is often considered more damaging to *A. mellifera* colonies than the more notorious *V. destructor* (Anderson and Morgan, 2007; Buawangpong *et al*., 2015; de Guzman *et al*., 2017). More recently in 2008, a second *Varroa* species, *V. jacobsoni*,was found to have also shifted hosts from *A. cerana* to *A. mellifera* in Papua New Guinea (PNG) (Roberts *et al*., 2015; Anderson, 2008). *V. jacobsoni* is a native parasite of *A. cerana* in Southeast Asia, which has invasively spread to the Pacific region. Since its first detection in 2008, *V. jacobsoni* has caused significant colony losses in PNG but remains restricted to this population. Around the same time, *T. mercedesae* invasively spread across PNG from neighbouring Indonesia (West Papua) also causing significant harm to *A. mellifera* colonies. Beekeeping in PNG is a cottage industry that has seen significant declines with current annual honey production at ca. 35 tonnes compared to previous peak production levels of more than 100 tonnes per annum (Orlegge and Gonapa, 2011). The PNG environment is well-suited to honey production and there is high domestic and international demand for quality honey. However, industry growth requires effective management strategies for these serious mite pests in PNG, which also present significant biosecurity threats to the region.

Globally, management strategies for *Varroa* and *Tropilaelaps* mites are heavily reliant on synthetic and organic chemical treatments (Rosenkranz *et al*., 2010). These chemicals are all developed and assessed for *V. destructor* in temperate environments. While likely effective against *T. mercedesae* and *V. jacobsoni*, there are important biological and environmental factors that may reduce their effectiveness in PNG. From the few studies on *T. mercedesae* in Asia, formic acid and the commercial products Bayvarol^®^ (flumethrin), Apistan^®^ (tau-fluvalinate) and Check-Mite^®^ (coumaphos) can offer effective control (Kongpitak *et al*., 2008; Pettis *et al*., 2017; Wilde *et al*., 2000; Mahmood *et al*., 2011). However there is no literature on the effectiveness of any chemicals against the new *V. jacobsoni*, although Bayvarol^®^ has started being used by some beekeepers in PNG. Other concerns with the use of chemicals are the negative effects this can have on colony health (Mullin *et al*., 2010; Rangel and Tarpy, 2015; Collins *et al*., 2004), contamination of hive products (Calatayud-Vernich *et al*., 2018; Wilmart *et al*., 2016), and risk of mites developing chemical resistance, as has occurred for *V. destructor* (Sammataro *et al*., 2005). These issues are likely to be exacerbated in PNG, given the complexities of providing training and educational services on the correct use of chemical controls to a dispersed population of rural beekeepers. The cost of chemical treatment is also prohibitive for many smallholder beekeepers, commonly resulting in under-treatment of colonies or no treatment at all.

Cultural control strategies may offer low-cost alternatives for smallholder beekeepers in PNG. Both *Varroa* and *Tropilaelaps* mites feed and reproduce on the developing brood of *A. mellifera*, hence an interruption of brood rearing to remove the supply of food can effectively reduce mite populations. This approach is likely to be particularly effective against *T. mercedesae*, as they are unable to feed on adult bees and can only survive several days without access to brood (Woyke, 1984; Khongphinitbunjong *et al*., 2019), whereas *V. jacobsoni* can feed and survive longer on adult bees (Boot *et al*., 1993; Calatayud and Verdú, 1994). Techniques for interrupting brood rearing include caging the queen to prevent egg-laying and/or the removal of brood (Khongphinitbunjong *et al*., 2014; Pettis *et al*., 2017; Wilde *et al*., 2000; Woyke, 1985). However, as these strategies are more labour-intensive and sometimes destructive of brood, they are generally not seen as practical for beekeepers. Queen caging techniques have been demonstrated experimentally with high effectiveness against *T. mercedesae* in Afghanistan and Vietnam (Woyke, 1984; Woyke, 1985). Pettis *et al* (2017) showed that splitting a hive into a parent and nucleus colony was also effective against *T. mercedesae* in Thailand, despite not creating a complete break in brood rearing. However, the effectiveness and suitability of these types of strategies against both *V. jacobsoni* and *T. mercedesae* in the PNG context is unknown.

Here we provide the first data on the comparative effectiveness of Bayvarol^®^ and cultural control strategies against *V. jacobsoni* and *T. mercedesae* in PNG. We found Bayvarol^®^ was highly effective against *V. jacobsoni*, but less effective against *T. mercedesae*. However, cultural control strategies using queen caging and queen removal techniques were very effective for *T. mercedesae*. These findings highlight clear potential for developing an integrated approach to mite control in PNG and provide valuable biosecurity information to the region.

## Methods

### Study site

The PNG Highlands has a temperate climate that is well suited to honey production with *A. mellifera*. Colonies produce brood year round although the dry season (May to September) is typically a period of lower brood production. The experiment was conducted in the Lufa District of the Eastern Highlands Province (EHP) starting in May 2018 using 20 colonies at site 1 (GPS, 6° 15’ 15” S 145° 26’ 6” E) and eight colonies at site 2 (GPS 6° 14’ 56” S 145° 26’ 5” E) to allow seven replicate hives per treatment. All colonies had not received any mite management for more than six months, were free of brood disease and of comparable strength in terms of brood and bee population.

### Experimental design

Experimental hives were randomly assigned to a treatment or control group with seven replicates per treatment (see Table 1). Colonies receiving a Bayvarol^®^ treatment were given the recommended dosage rates of two flumethrin-impregnated strips per five frames of brood and removed after eight weeks. The queen caging treatment was adopted from Woyke (1984) where queens were caged within the colony to prevent egg laying and released after 28 days. This created a brood-free period of ca. three days after all existing brood emerged. The artificial swarm treatment also aimed to create a brood-free period of at least three days in both the parent and new colony. This involved transferring the queen from the parent colony along with adult bees shaken from frames into a new hive box with only foundation frames, so as to mimic swarming. The entrance of the new colony was blocked for 24 hours to reduce bees returning back to the parent colony. Unfortunately, this was not done effectively and many bees returned to the parent colonies in the apiary. As a result, the new colonies had small starting populations and struggled to establish during the experiment. The data collected from these colonies was considered unreliable and therefore excluded from the analysis. In theory, the new ‘swarm’ colony is prevented from producing brood for several days until the bees can create brood comb for the queen. The parent colonies were left to raise a new queen, during which no eggs are laid for at least 28 days. Control colonies were left untreated for mite during the experiment. Initial levels of *V. jacobsoni* and *T. mercedesae* were estimated using 24 hour natural mite fall onto a sticky board (Mann Lake Ltd. Minnesota, USA) followed by additional estimates using sugar-shake, brood uncapping and frame bump techniques (described below) for comparison. Mite levels were monitored monthly mite with sticky boards until September 2018, where a final estimate was made again using all monitoring methods.

**Table 1.**
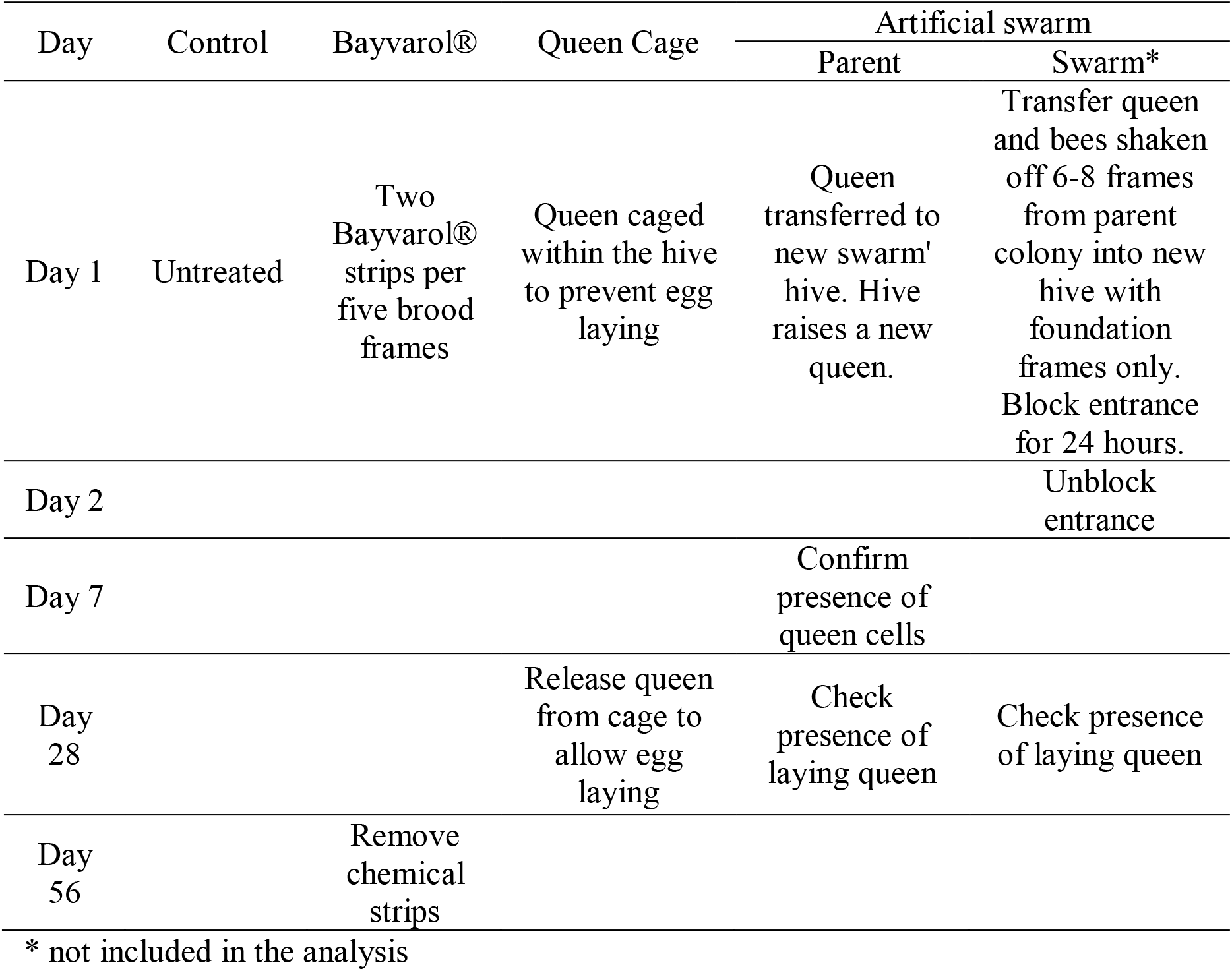
Mite management experiment comparing Bayvarol^®^ with two brood interruption strategies against *V. jacobsoni* and *T. mercedesae* in PNG.

| Day | Control | Bayvarol® | Queen Cage | Artificial swarm |  |
| --- | --- | --- | --- | --- | --- |
|  |  |  |  | Parent | Swarm* |
| Day 1 | Untreated | Two Bayvarol® strips per five brood frames | Queen caged within the hive to prevent egg laying | Queen transferred to new swarm' hive. Hive raises a new queen. | Transfer queen and bees shaken off 6-8 frames from parent colony into new hive with foundation frames only. Block entrance for 24 hours. |
| Day 2 |  |  |  |  | Unblock entrance |
| Day 7 |  |  |  | Confirm presence of queen cells |  |
| Day 28 |  |  | Release queen from cage to allow egg laying | Check presence of laying queen | Check presence of laying queen |
| Day 56 |  | Remove chemical strips |  |  |  |

\* not included in the analysis

### Monitoring methods

Several mite monitoring methods were used in this study. Most are designed for *V. destructor*, although there has also been some assessment against *T. mercedesae* (Pettis *et al*., 2013; Anderson and Roberts, 2013). Sticky boards were the main monitoring method used as they were expected to give the most reliable estimates of both mite species and offer less technical variation between users. This method involved placing a sticky board on the bottom of each experimental hive for 24-hours and then counting fallen mites under low magnification to estimate colony mite populations. The sugar-shake method is well established for *Varroa* mites but considered less reliable for *T. mercedesae* as they have shorter phoretic periods. This method involved collecting ca. 300 adult bees from the brood nest into a container with a mesh lid and adding powdered sugar to coat the bees. The container and bees were then shaken over a tray to dislodge and count phoretic mites. Uncapping 100 worker brood cells using forceps was also used and considered an effective method for detecting mites, particularly *T. mercedesae* as a high percentage of the mite population is present inside sealed brood cells. In addition, we used the ‘bump test’ following Pettis et al. (2013). After cell uncapping, the frame was firmly bumped twice over a tray to dislodge and count mites.

### Statistical analysis

*V. jacobsoni* and *T. mercedesae* numbers were estimated from the average number of mites counted from three rows of the sticky board. Suitability of this estimation method was cross-checked and confirmed by counting the total mites counted on several sticky boards. Mite treatment data was analysed in GraphPad Prism 7 using two-way ANOVAs to compare changes in log transformed mean mite numbers across treatments between June and October. Colonies with initial mite numbers less than 10 were excluded from the analysis, although this still allowed at least four replicates per treatment. Efficacy of each treatment at each time point was calculated using the geometric mean mite numbers and the equation; efficacy = (untreated mean – treated mean)/untreated mean x 100 (Kongpitak *et al*., 2008).

## Results

### *Efficacy of mite management strategies against* V. jacobsoni *and* T. mercedesae

For *V. jacobsoni*, two-way ANOVA showed a significant main effect for treatment (*F*_(3,84)_ = 15.73, *p* < 0.0001), but not for month (*F*_(4,84)_ = 2.32, *p* = 0.064) or any interaction (*F*_(12,84)_ = 1.062, *p* = 0.403). Tukey HSD tests revealed that mean *V. jacobsoni* numbers were significantly reduced only for the Bayvarol^®^ treatment compared to the untreated control hives and remained significantly low through to September (Figure 1). The brood manipulation strategies using queen caging or creating an artificial swarm showed minimal effect on *V. jacobsoni* levels, despite indications of mite suppression in July.

**Figure 1.**
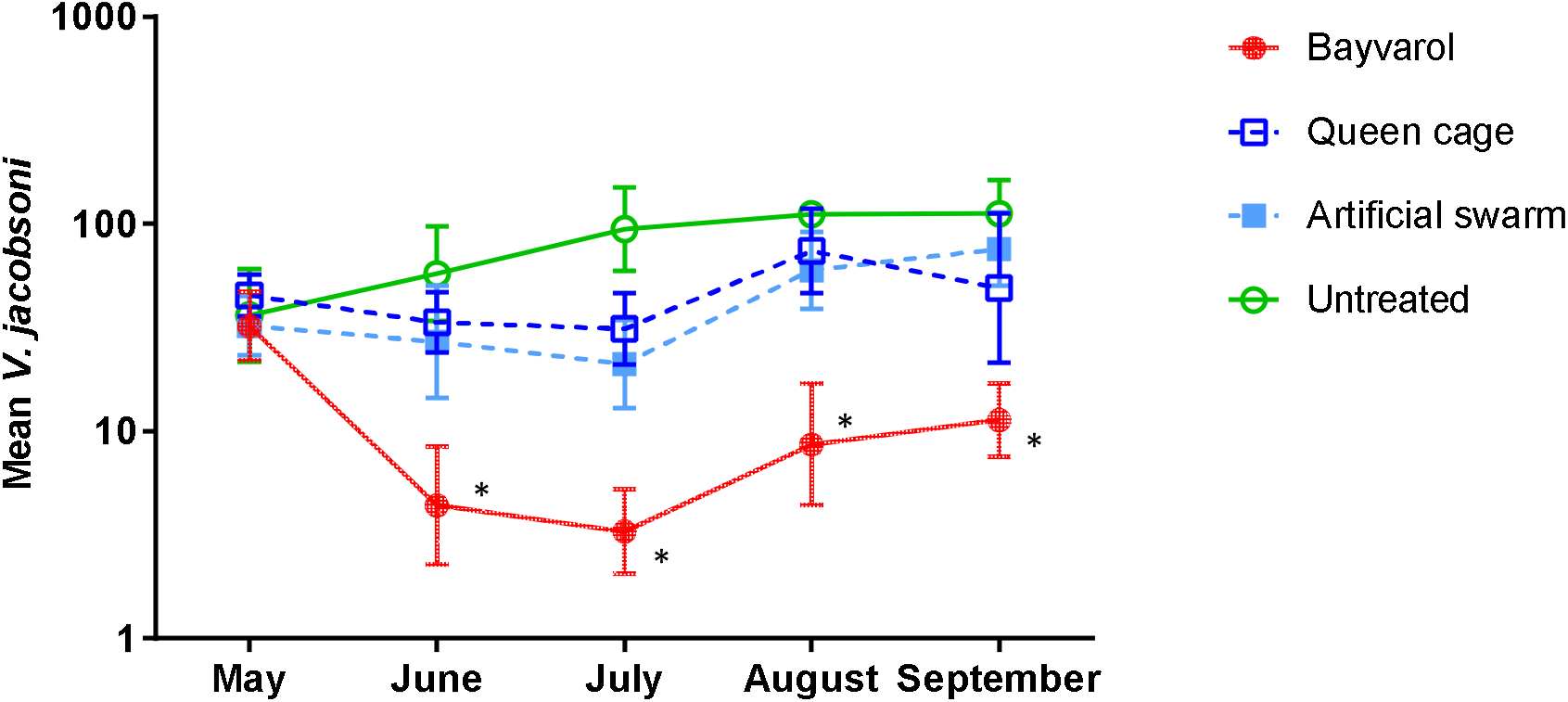
Mean numbers of *Varroa jacobsoni* estimated from sticky boards in hives before (May) and after treatment (June to September). * denotes a significant difference (P < 0.05) from the untreated control.

For *T. mercedesae*, two-way ANOVA showed a significant main effect for treatment (*F*_(3,105)_ = 4.78, *p* = 0.004), month (*F*_(4,105)_ = 10.65, *p* < 0.0001) and a significant interaction (*F*_(12,105)_ = 2.41, *p* = 0.008). Tukey HSD tests revealed that mean *T. mercedesae* numbers were significantly reduced by the queen caging treatment in June and July, and the queen removal treatment by July compared to the untreated control hives (Figure 2). Bayvarol^®^ did not significantly reduce *T. mercedesae* numbers compared to untreated hives, although there was a trend for lower numbers.

**Figure 2.**
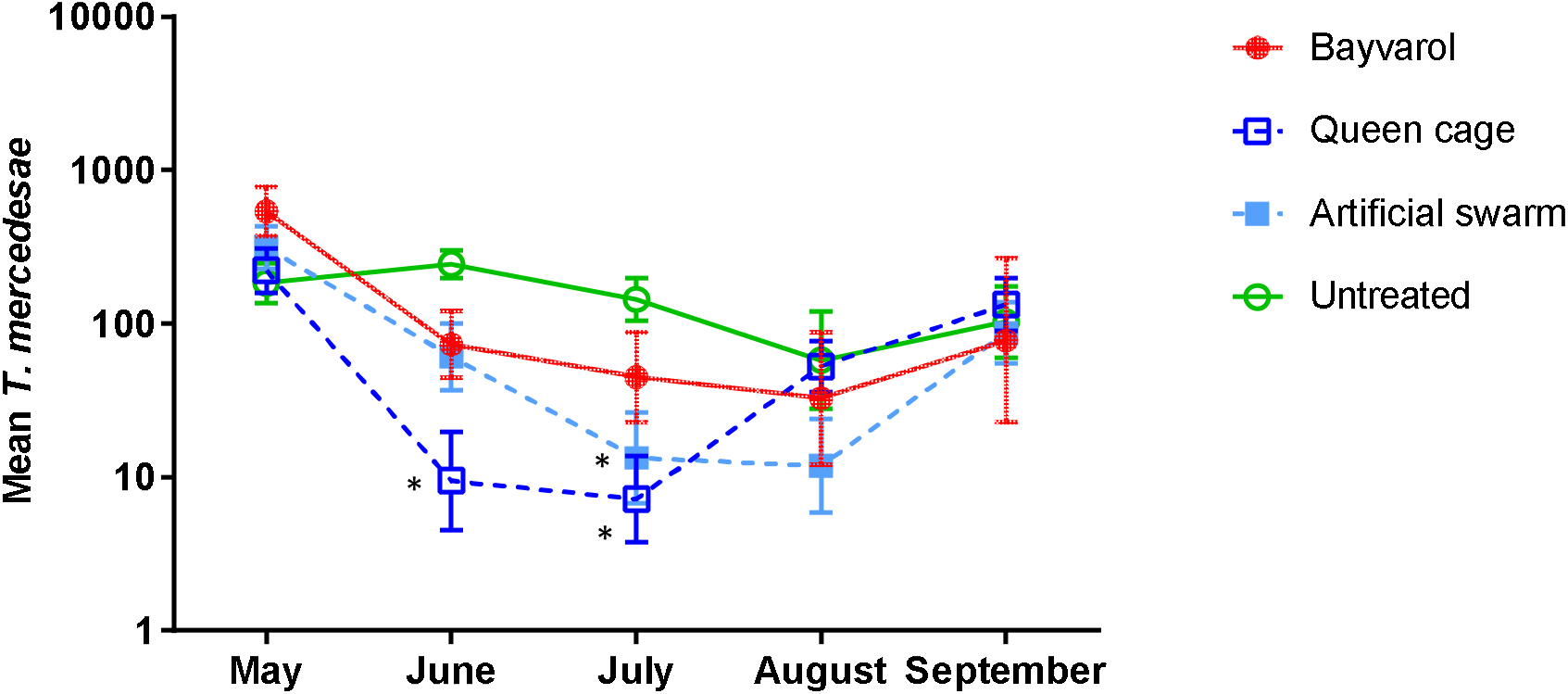
Mean numbers of *Tropilaelaps mercedesae* estimated from sticky boards in hives before (May) and after treatment (June to September). * denotes a significant difference (P < 0.05) from the untreated control.

The calculated efficacy of each treatment at each time point is shown in Table 2. The Bayvarol^®^ treatment was the most effective control strategy for *V. jacobsoni*, maintaining high efficacy (>90%) through to September. In contrast, Bayvarol^®^ was not very effective against *T. mercedesae* as a knock-down treatment, only reaching ca. 70% efficacy in June and July before rapidly losing efficacy. The brood manipulation strategies both had high efficacy (> 90%) against *T. mercedesae* early, but this was not sustained for more than two months. And while not significant for *V. jacobsoni*, there was 68% and 79% efficacy in July for the queen caging and artificial swarm treatment respectively.

**Table 2.** Percentage efficacy of mite treatments for *V. jacobsoni* and *T. mercedesae*.

|  | *Varroa jacobsoni* |  |  | *Tropilaelaps mercedesae* |  |  |
| --- | --- | --- | --- | --- | --- | --- |
|  | Bayvarol® | Queen cage | Swarm (Parent) | Bayvarol® | Queen cage | Swarm (Parent) |
| June | 93 % | 42 % | 56 % | 71 % | 96 % | 76 % |
| July | 97 % | 68 % | 79 % | 70 % | 95 % | 91 % |
| August | 93 % | 35 % | 47 % | 44 % | 11 % | 81 % |
| September | 91 % | 56 % | 33 % | 23 % | 0 % | 15 % |

### Comparison of mite monitoring methods

Spearman non-parametric correlations between the different methods for estimating mite numbers found the sticky board and all three other methods (sugar-shake, uncapping and bump test) were significantly correlated (P < 0.05) for both *V. jacobsoni* and *T. mercedesae* (Figure 3, Table 3). While these methods clearly differ in their sensitivity to detect low level mite infestations, this finding suggests that these other methods are still informative monitoring options for smallholders and are likely more affordable.

**Figure 3.**
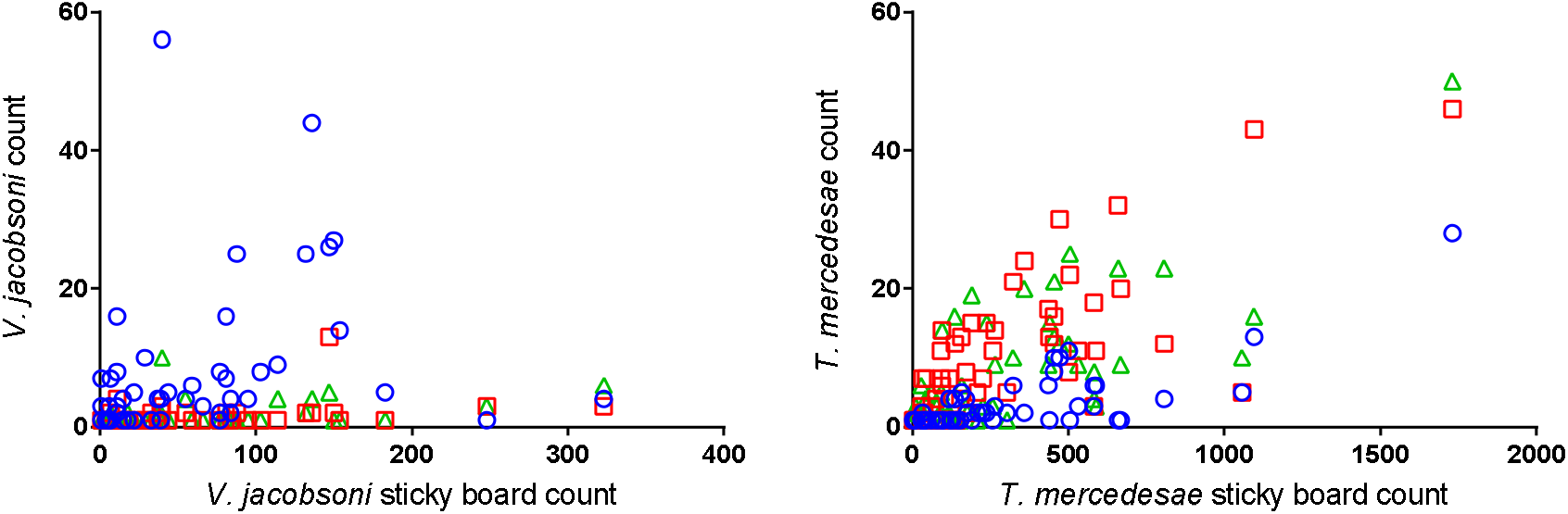
Correlation between mite numbers estimated from the sticky mat and sugar shake (circles), uncapping (square) and bump (triangles) monitoring methods

**Table 3.**
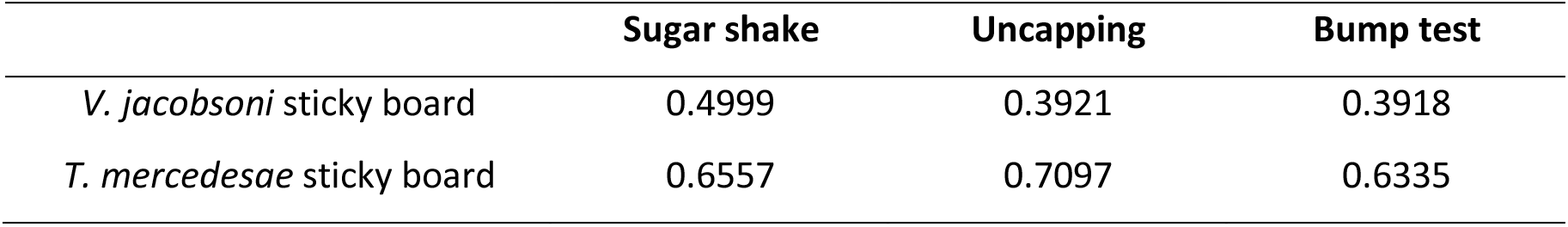
Spearman non-parametric correlation r values showing significant positive relationships for mite numbers estimated from sticky mats and sugar-shake, uncapping and bump test methods.

## Discussion

This study has provided the first comparative data for different management strategies against *V. jacobsoni* and *T. mercedesae* in PNG. This information is vital for the development and adoption of effective ‘best practice’ guidelines to control mite populations that are suitable for PNG beekeepers. Currently, Bayvarol^®^ is the only chemical available in PNG and our findings support its continued use to effectively reduce *V. jacobsoni* populations for several months. However, Bayvarol^®^ did not perform well against *T. mercedesae* under these study conditions. This is likely explained by the different phoretic behaviours of the two mites, with *T. mercedesae* spending much less time on adult bees where they can be exposed to contact chemicals. This was also suggested as a reason for the low efficacy of amitraz, another contact chemical, against *T. mercedesae* in Thailand (Pettis *et al*., 2017). However, another study in Thailand report 100% efficacy from three common *Varroa* contact chemicals (flumethrin, coumaphos and tau-fluvalinate) (Kongpitak *et al*., 2008). It is likely that colony size and brood levels can heavily influence the effectiveness of contact chemicals and can be better optimized for PNG beekeepers. For example, consolidating the brood to a single box with a queen excluder, which is not commonly practiced in PNG or applied in our study, may offer more consistent treatment against both mite species.

Reliance on a single chemical presents a high risk for mite resistance to develop in PNG. Particularly when there are local reports of chemical misuse such as undertreating to reduce costs or leaving in treatment strips past the recommended period (J. Jave pers. comm, 20^th^ May 2018). Rotation with other chemical and cultural management strategies must be a priority for the PNG beekeeping industry. Organic chemicals may be useful alternatives compared with synthetic products such as coumaphos and amitriz, as they have a lower risk of residues in bee products and of mite resistance developing (Rosenkranz *et al*., 2010). Formic acid may be a good option as it is the only available product that can penetrate capped brood cells to reach mites (Fries, 1991). It has been shown to give good control of both *V. destructor* and *T. mercedesae* elsewhere (Mahmood *et al*., 2011; Pettis *et al*., 2017; Rosenkranz *et al*., 2010), although results can be variable under different environmental conditions. Pettis et al (2017) found both Hopguard^®^ and sulfur also significantly reduced *T. mercedesae* populations, although the dose and timing of application was critical to limit negative impact on adult bees and brood. Mite populations also recovered after two months using these products, which is equivalent to the results of the cultural control strategies tested in this study. A suitable starting point for beekeepers in PNG may be to assess the effectiveness and practical suitability of formic acid in the commercially available sustained release Mite-Away Quick Strips^®^, in order to reduce application variability and health and safety risks.

Cultural control strategies aimed at brood manipulation have clear potential to effectively manage mite populations as part of an integrated approach. These strategies are often not used because they are seen as too labour intensive however they may be well suited to PNG beekeepers who typically have fewer colonies and the cost of chemical treatments is likely a more limiting factor. Our results demonstrate two cultural control methods that were very effective against *T. mercedesae*, which can be further optimised for PNG beekeepers. The ability of *V. jacobsoni* to survive on adult bees for weeks without brood clearly limits the effectiveness of brood interruption strategies. However, there was slower population growth under these strategies compared to the untreated hives, which may prove more effective during peak mite population increases. Extending the effects for *T. mercedesae* may also be possible. The fast reproductive rate of *T. mercedesae* can facilitate a rapid recovery of mite numbers, although reintroduction of mites from neighbouring untreated hives may have had a greater influence in this study. In practice, a mite treatment would be applied to all hives in an apiary to minimise the rate of reintroduction, resulting in longer term control. We are also confident the artificial swarm treatment can offer further benefits to beekeepers if better executed. Another ‘swarm’ colony made in parallel to the experiment and moved several kilometers from the study sites was well established four months later and a sugar-shake in September found relatively high *V. jacobsoni* numbers (50 mites) but no *T. mercedesae*.

The productivity and growth of the beekeeping industry in PNG requires integrated mite management protocols that are based on the best available information to inform management decisions, extension and training for smallholder beekeepers. Research for best practice management in PNG and throughout the Asia-Pacific region needs to take into consideration the effectives of the treatment, but also the risks for beekeepers in up taking new approaches, issues of access and associated costs and difficulty of implementation. These approaches need to be best optimized to suit the local conditions to enable access and enhance uptake among smallholder beekeepers. This study has demonstrated that cultural controls can reduce mite levels for both *V. jacobsoni* and *T. mercedesae*, however further optimization is required to improve efficiencies for both species and provide ‘best practice’ guidelines. These methods do not require high capital investment and can help to improve rates of uptake for rural beekeepers and reduce colony losses and dependencies on chemical controls. Further focus on pest and disease management solutions that are appropriate for rural beekeepers are required to improve the productivity, profitability and sustainability of the apiculture sector throughout PNG and the Asia-Pacific region.

## Acknowledgments

The authors wish to appreciate the dedication time, guidance and invaluable technical contributions of the Apicultural Officers of National Department of Agriculture and Livestock apiculture team and Coffee Industry Cooperation, namely, Mr Jonah Buka, Mr George Waenavi, Mr Joachim Waugla, Mr Jesse Yawane, Mr Johna Neghia and Mr Shayne Loie Tumae.

## Notes

**Disclosure statement:** No potential conflicts of interest are reported by the authors.

**Funding details:** This work was supported by the Australian Centre for International Agricultural Research (ACIAR) under Grant LS/2017/100.

